# An analog to digital converter creates nuclear localization pulses in yeast calcium signaling

**DOI:** 10.1101/357939

**Authors:** Ian S Hsu, Bob Strome, Sergey Plotnikov, Alan M Moses

**Affiliations:** Department Of Cell & Systems Biology University of Toronto

**Keywords:** stochastic pulsing transcription factor, Crz1, time delay model, mathematical modeling, calcineurin pathway

## Abstract

Several examples of transcription factors that show stochastic, unsynchronized pulses of nuclear localization have been described. Here we show that under constant calcium stress, nuclear localization pulses of the transcription factor Crz1 follow stochastic variations in cytoplasmic calcium concentration. We find that the size of the stochastic calcium pulses is positively correlated with the number of subsequent Crz1 pulses. Based on our observations, we propose a simple stochastic model of how the signaling pathway converts a constant external calcium concentration into a digital number of Crz1 pulses in the nucleus, due to the time delay from nuclear transport and the stochastic decoherence of individual Crz1 molecule dynamics. We find support for several additional predictions of the model and conclude that stochastic input to nuclear transport may produce digital responses to analog signals in other signaling systems.

## Introduction

Cells transmit information through signaling pathways. Rather than simple “ON” or “OFF” responses, several key pathways (p53, NF-icB, and others) are now appreciated to encode information in the dynamics of the signaling response(1-9). Here we focus on the calcium signaling pathway in yeast, which controls gene transcription through frequency modulation (FM) of the transcription factor Crz1(9). In this system, the analog external calcium concentration is converted into the frequency of digital pulses of nuclear localization (discrete rapid rising and falling of nuclear concentration on the order a few minutes).

Mechanistic models of FM pulsatile transcription factors have relied on negative feedback coupled with positive feedback(7,8,10,11) or delayed negative feedback(11,12). However, whether there is a negative feedback loop in calcium signaling pathway that can generate Crz1 pulses is unclear (discussed further below). Furthermore, in each cell, during each Crz1 nuclear localization pulse, approximately 500 Crz1 molecules are transported in and out of the nucleus in a coordinated fashion(13), but these pulses are stochastic (and not synchronized between cells). To our knowledge, no mechanistic model of this process has yet been proposed.

One possibility is that Crz1 nuclear localization pulses are connected to variation in cytoplasmic calcium concentration ([Ca^2^+]_cyt_) since calcineurin is activated by calcium(14). Calcium pulses have been observed in many cell types(15-19), but the connections between cytoplasmic calcium concentration and Crz1 localization have not been analyzed in single cells. Mechanistic models of Crz1 regulation through calcium signaling do not predict external calcium concentration ([Ca^2^+]_ext_)-induced oscillation of [Ca^2^+]_cyt_(20). Further, although average Crz1 nuclear localization increases when [Ca^2+^]ext increases(9), [Ca^2+^]cyt is known to be under tight homeostatic control: the average [Ca^2+^]cyt remains similar under a wide range of [Ca^2^+]ext(21,22). Variation in [Ca^2+^]cyt is unlikely to follow the frequency of Crz1 pulsatility, which increases when [Ca^2+^]ext increases(9). Thus, the relationship between calcium and Crz1 pulses remains unclear.

In this study, we examined the connection between [Ca^2+^]cyt and Crz1 pulsatile dynamics through dual fluorescence time-lapse microscopy(23). We found that cytoplasmic calcium concentration varies stochastically at the single cell level, showing pulses on the timescale of 10-100 seconds. We observed overshoots of the calcium concentration, strongly implicating calcium channels in these pulses. We found that Crz1 pulses tend to follow these calcium pulses, but that the relationship is not simple: multiple Crz1 pulses may follow each calcium pulse, and the number of Crz1 pulses depends on the size of the calcium pulse. We modulated calcium channel activity and found much larger calcium pulses, which led to greater numbers of Crz1 pulses. To explain how [Ca^2^+]_cyt_ affects Crz1 nuclear localization, we developed a stochastic model of Crz1 nuclear localization and tested predictions in the experimental data. In general, stochastic pulses in signaling dynamics may be generated by time-delayed responses to fluctuations in second messenger concentration.

## Results

### Calcium pulses are observed when yeast cells are under calcium stress

In order to study the relationship between the dynamics of [Ca^2^+]_cyt_ and Crz1 nuclear localization, we constructed a dual Crz1-Calcium reporter strain and measured dynamics using time-lapse microscopy. We tagged Crz1 with the mCherry fluorescent protein in a strain with a cytoplasmicly expressed calcium sensor GCaMP3(16), and recorded movies on a confocal fluorescence microscope (see Methods). As in previous studies(9), we observed stochastic and rapid increases and decreases of [Ca^2^+]_cyt_ when yeast are under calcium stress (Figure 1 A, supplementary video 1). We noticed that these “calcium pulses” (defined by a threshold ratio above background, see Methods) are followed by an overshoot of calcium concentration below the resting level (average of 50 largest calcium local maxima in a representative time lapse movie is shown in Figure 1B left panel), whose depth is positively correlated to the height of the pulse (Figure 1B, right panel, R^2^= 0.37). This overshoot cannot fit an exponential curve and suggests negative feedback on [Ca^2^+]_cyt_, which is consistent with the predictions of calcium models constructed in previous studies of homeostasis(24,25).

**Figure 1.**
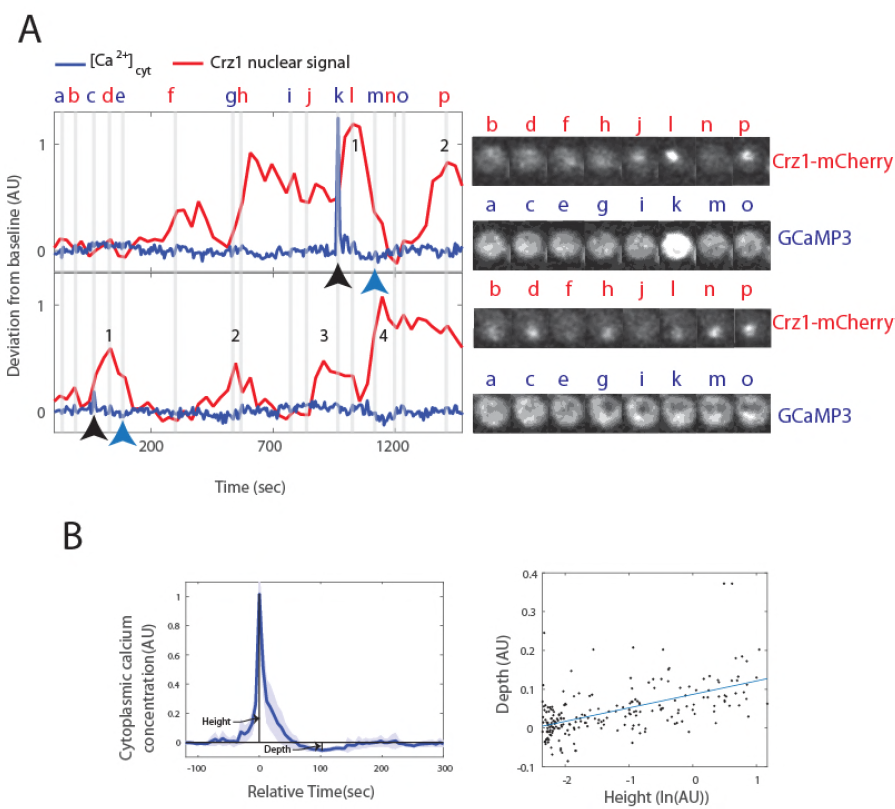
Calcium pulses with overshoots are found in yeast cells under calcium stress. A) Left panel shows examples of single cell trajectories for Crz1 (red trace) and calcium (blue trace) and snapshots at representative time points from two cells close to each other in the original field of view. Numbers along the red trace indicate the Crz1 pulses identified following a calcium pulse (black arrow below blue trace). Blue arrow indicates the local minimum (so-called overshoot) following the calcium pulse. Images in the right panel show the mCherry channel and GCaMP3 channel at points indicated in the left panel. B) Left panel shows the average trace of 50 calcium pulses. Shaded area shows 95% CI of the average trace. Maxima of calcium pulses are aligned to time = 0 sec (Relative time). An overshoot can be found around time = 100 sec. In the right panel, each dot represents a single calcium pulse. The x-axis is in natural log of peak height, while the y-axis is the depth of the overshoot. Blue line shows a linear fit (R^2^ = 0.37).

### Quantification of calcium pulses and Crzl pulses suggests an analog to digital converter

Individual cell trajectories do not show a simple relationship between [Ca^2+^]cyt and Crz1 nuclear localization (Figure 1A), so we next sought to understand how Crz1 pulses (see Methods for definition of Crz1 pulses) are affected by calcium pulses. We analyzed the distribution of the time differences between a calcium pulse and following Crz1 pulse(s) (using so-called pulse-triggered averaging(23)). The coherence of the first and second pulses suggested to us that one or more Crz1 pulses follows a single calcium pulse (Figure 2A). We therefore compared the time until the first Crz1 pulse within 10 minutes of a calcium pulse to the time until the first Crz1 pulse within 10 minutes from a randomly chosen cell that may or may not contain a calcium pulse. Consistent with our hypothesis, we found that the time until the first Crz1 pulses after calcium pulses shows reduced standard deviation (122.47 seconds vs. 161.25 seconds, F-test, p<0.005, n = 168) and occurs sooner than observed in randomly chosen cells (83.01+-9.58 seconds vs. 210.62+-28.89 for random, two-tailed t-test, p <10^−15^, n = 193). A similar range of time differences is also observed in cross correlation analysis (supplementary figure 1). Furthermore, the distributions of time differences disperse when the order of Crz1 pulses increases (Figure 2A, e.g., 122.47 seconds, n = 81 for first pulses vs. 181.66 seconds, n = 45 for second pulses, F-test, p<0.005), suggesting that the effect of the calcium pulse on Crz1 dynamics decreases over time. These results suggest that one calcium pulse can lead to multiple Crz1 pulses.

**Figure 2.**
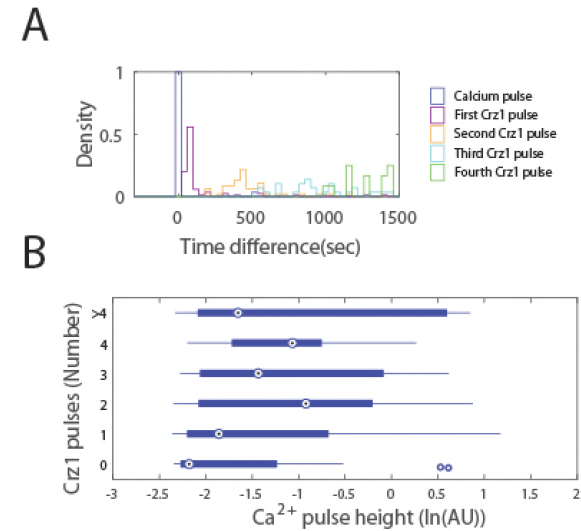
A calcium pulse is followed by multiple Crz1 pulses, and the number of Crz1 pulses is positively correlated to the height of the calcium pulse. A) The density of first, second third and fourth Crz1 pulses (purple, yellow, cyan and green, respectively) is plotted as a function of the time they occur after the calcium pulse (blue). B) Sample sizes of each number of Crz1 pulses are 29 for 0, 73 for 1, 43 for 2, 21 for 3, 16 for >3. Blue boxes indicate the 25% − 75% range, large circles represent the mean, lines represent the range of the data, and individual points show the locations of two outliers.

We sought to identify factors that determine the number of Crz1 pulses that follow each calcium pulse. We found that the number of Crz1 pulses after a calcium pulse is positively correlated to the height of that calcium pulse (Figure 2B, generalized linear model regression with Poisson distribution, slope = 0.16+-0.10). To test whether a calcium pulse causes Crz1 pulses, we tested whether there is a correlation between calcium pulse height and the number of Crz1 pulses before a calcium pulse. If an (unmeasured) third factor affects both the height of a calcium pulse and the number of the Crz1 pulses in a cell, we expect more Crz1 pulses both before and after large calcium pulses. We found that the number before is not correlated with calcium pulse size (slope = −0.10+-0.13, generalized linear model regression with Poisson distribution). This result is consistent with the idea that calcium pulses can cause more than one Crz1 pulse, and suggests that calcium pulse heights are converted into digital numbers of Crz1 pulses.

### Artificially increased calcium pulse height supports the analog to digital converter model

The hypothesis of an analog to digital converter between calcium pulse height and Crz1 pulse number predicts that the average number of Crz1 pulses following calcium pulses could be made larger by artificially inducing larger calcium pulses. To test this prediction, we treated cells with nifedipine (See Methods). By doing so, we reliably induced synchronized calcium pulses (supplementary video 2) that were on average twice as large as the stochastic pulses observed at steady state in 0.2M calcium treatment alone (Figure 3A, compared to Figure 1B). The majority of these large calcium pulses are followed by at least 4 Crz1 pulses that disperse over time (Figure 3B). As predicted by the model, the number of Crz1 pulses after a calcium pulse is positively correlated to calcium pulse height, and the range is larger than untreated range (Figure 3C, compared to Figure 2B, generalized linear model regression with Poisson distribution, slope = 0.37+-0.11). In individual cells, nuclear Crz1 now clearly appears to oscillate while no oscillations are observed in cytoplasmic calcium concentration (supplementary figure 2). These results confirm that single calcium pulses are followed by multiple Crz1 pulses, and support the idea that an analog-to-digital converter in the calmodulin/calcineurin signaling pathway converts calcium pulse height into Crz1 pulse number.

**Figure 3.**
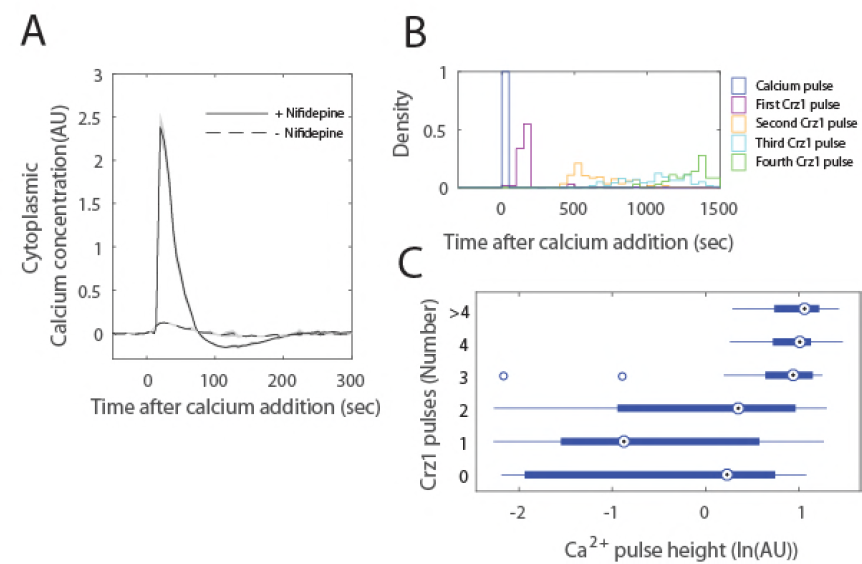
Artificially induced calcium pulses with large average height are followed by more Crz1 pulses. A) Average single cell trajectory (without aligning the maxima) after calcium addition shows synchronized calcium pulses in cells treated with nifedipine (solid line is mean GCaMP3 signal), not in untreated cells (broken line is the mean GCaMP3 from untreated cells. Shaded areas show the 95% CI of mean. B) The density of first, second third and fourth Crz1 pulses (purple, yellow, cyan and green, respectively) is plotted as a function of the time they occur after the calcium pulse (blue). C) Sample sizes of each number of Crz1 pulses are 16 for 0, 36 for 1, 36 for 2, 30 for 3, 48 for 4, 14 for >4. Blue boxes indicate the 25% − 75% range, large circles represent the mean, lines represent the range of the data, and individual points show the locations of outliers.

### A simple time delay model can reproduce the properties of Crzl pulses after a calcium pulse

To explain the mechanism of the analog to digital converter, we considered a two-step process in single cells (Figure 4A). The first step is that external calcium concentration leads to cytoplasmic calcium pulses (4A, blue trace) through stochastic channel opening, and the second step is that a calcium pulse leads to nuclear Crz1 pulses (4A, red trace) through the calcineurin pathway.

**Figure 4.**
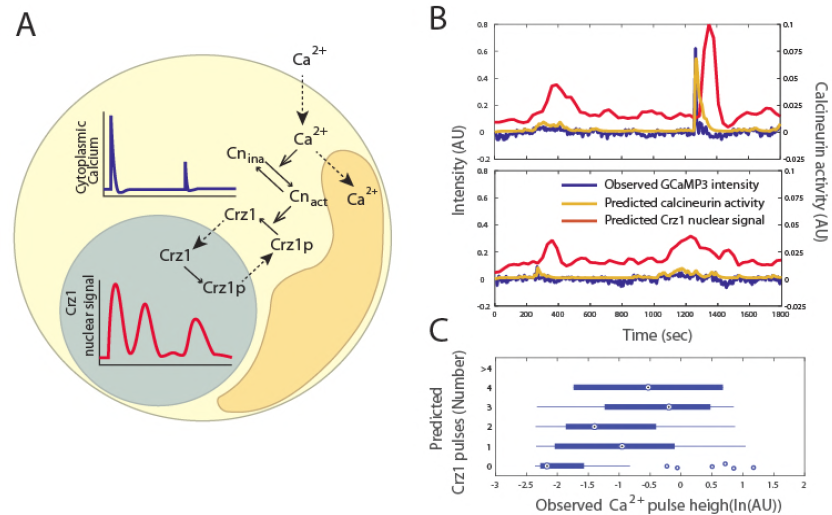
A time delay model for Crz1 nuclear pulsing can qualitatively reproduce Crz1 dynamics after calcium pulses. A) In this single cell model, Ca^2^+ in the cytoplasm (yellow shaded area) is controlled through a two-channel system (dotted arrows crossing from outside the cell to inside, and crossing from yellow shaded area to orange shaded area representing the vacuole). Calcineurin is activated (Cn_ina_ to Cn_act_) by cytoplasmic calcium, and leads to Crz1 dephosphorylation (Crz1p to Crz1). Dephosphorylated Crz1 is imported (dotted arrow) from cytoplasm (yellow shaded area) to nucleus (grey shaded area) and phosphorylated Crz1 is exported (dotted arrow from nucleus to cytoplasm). B) The Crz1 pulses produced by the model (red trace) following calcium pulses (blue trace). Gold trace shows the calcineurin activity predicted by the model. C) Blue boxes indicate the 25% − 75% range, large circles represent the mean, lines represent the range of the data, and individual points show the locations of outliers. Sample sizes of each number of Crz1 pulses are 68 for 0, 72 for 1, 31 for 2, 9 for 3, 2 for 4, 0 for

A negative feedback loop in the calcineurin pathway could lead to oscillation of calcineurin activity and drive Crz1 pulses, but we decided not to include one in our model for two reasons. First, the known negative feedback loop in the calcineurin pathway through Rcn1 does not appear to affect Crz1 pulsatility. Rcn1 is an inhibitor of calcineurin that is degraded when phosphorylated and is dephosphorylated by activated calcineurin (26,27), thus leading to negative feedback. However, the negative feedback loop is thought to be controlled by the protein abundance of Rcn1, as phosphorylation does not prevent Rcn1’s inhibition of calcineurin(26). Since Crz1 pulsatility occurs when protein synthesis is inhibited by cycloheximide (supplementary figure 3) we consider it unlikely that Rcn1 provides negative feedback through changes in protein abundance. Second, models with feedback mechanisms seem incompatible with the observation that the frequency but not the amplitude of Crz1 pulses increases when the affinity of calcineurin docking site on Crz1 is enhanced(9). This is opposite to the expectation if Crz1 nucleocytoplasmic transition were driven by a feedback mechanism. Therefore, we worked toward models that do not include a feedback mechanism.

Previously, Crz1 nuclear localization dynamics were explained with a conformational switch model(22). This model assumes that the large number of phosphorylation sites on Crz1 leads to a sigmoid function relating calcineurin activity to Crz1 nuclear localization, so when calcineurin activity swings above and below a threshold, Crz1 sensitively reads out the perturbation in calcineurin activity and switches fully nuclear or cytoplasmic (22,28). This model would predict that calcium concentration crosses a threshold before each Crz1 pulse, and pulses stop once calcium oscillations decay below the threshold. However, calcium is not observed to pass a threshold before each Crz1 pulse in our data (supplementary figure 2D for examples).

We therefore considered another single cell model. Inspired by the observations on the population level that Crz1 pulses tend to occur within 100 seconds after calcium pulses and then disperse over time, and that larger calcium pulses lead to more Crz1 pulses, we constructed a discrete-time stochastic model that explains single cell Crz1 nuclear localization based on time delays during nuclear import and export with variation among Crz1 molecules. Time delay models have been constructed through different approaches, including deterministic and stochastic delay differential equations with a fixed or variable delay periods(12,29-31). We used a discrete-time Markov chain because it is simple to simulate trajectories. In the model, Crz1 molecules transit between the nucleus and the cytoplasm in a coordinated manner (show pulsing dynamics on average) only when calcineurin activity is very high due to a recent calcium pulse.As calcineurin activity slowly returns to its basal level, the coordinated transport of 500 Crz1 molecules in a single cell decoheres.

We model Crz1 nuclear signal by aggregating the states of individual Crz1 molecules after an increase in cytoplasmic calcium concentration of a single cell. As the input to the model, we provide calcium concentration, *Ca,* as a function of time, *t,* which can be obtained from experimental data (using GCaMP reporter fluorescence as a proxy).

We assume that calcineurin activity at time *t, Cn(t),* has an activation rate proportional to calcium concentration and a constant rate of decay. The discrete time dynamics of *Cn(t)* is described by

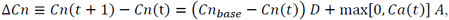

where *Cn_base_* is the basal activity of calcineurin, *D* is the decay rate of calcineurin activity, and *A* is the activation rate by cytoplasmic calcium. Although calcium concentration can never be negative, our GCaMP reporter data is normalized such that baseline fluorescence level is defined as 0. When calcium pulses overshoot, we obtain negative values, and hence include max[0, *Ca(t)]* in the equation above. The probability of a Crz1 molecule being imported is *Cn(t)* multiplied by the probability of dephosphorylation by an active calcineurin molecule (see Methods for details). Thus, when calcineurin activity increases, the probability of a Crz1 molecular being imported increases.

Once a Crz1 molecule is imported into the nucleus, it returns to the cytoplasm after it is phosphorylated in the nucleus, which we assume occurs at a constant rate. These chemical reactions can be formulated using a standard biochemical rate approach as

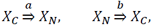

where *X_c_* and *X_N_* are cytoplasmic and nuclear Crz1 molecules, respectively, and *a* and *b* are the rates of delayed transports (denoted as thick arrows). In our Markov chain framework, we assume that transports are multistage, so the delay time follows a Gamma distribution with the two parameters related to the number of states and the transition probability (see Supplementary text for details).

This model can qualitatively reproduce Crz1 pulses after a calcium pulse (Figure 4B). The number of Crz1 pulses after a calcium pulse is positively correlated to the height of that calcium pulse (Figure 4C, generalized linear model regression with Poisson distribution, slope = 0.29+0.13), while the number of that before a calcium pulse is not (slope = 0.02+-0.16).

### Other predictions of the model are found in the experimental data

The time delay model also predicts other properties of Crz1 pulsatility. A first prediction is that the periodicity of a Crz1 trajectory is correlated with calcium pulse size, such that a Crz1 trajectory after a larger calcium pulse keeps oscillating longer. To test this prediction, we quantified the periodicity of Crz1 dynamics after the largest calcium pulse of each cell using a Gaussian Process model (see Methods), which computes the log-likelihood ratio (LLR) comparing a periodic to an aperiodic kernel. The LLR of post-calcium-pulse trajectories is correlated to the height of calcium pulses (Figure 5A). Calcium pulses larger than 0.11 show LLR significantly larger than that of the rest (two tailed t-test, sample sizes are 92 and 95, p < 10^-5^), which means larger calcium pulses lead to Crz1 dynamics that can be better described by a periodic Gaussian process. As a control, we also computed the LLR for pre-calcium-pulse trajectories, and found that the pre-calcium-pulse trajectories of for calcium pulse height above 0.11 are not statistically more periodic (p >0.1). Periodic dynamics after large calcium pulses would also be predicted by conformational switch model, where, after a larger calcium pulse, calcium oscillates longer and has more peaks crossing a threshold to trigger Crz1 pulse (supplementary figure 4).

**Figure 5.**
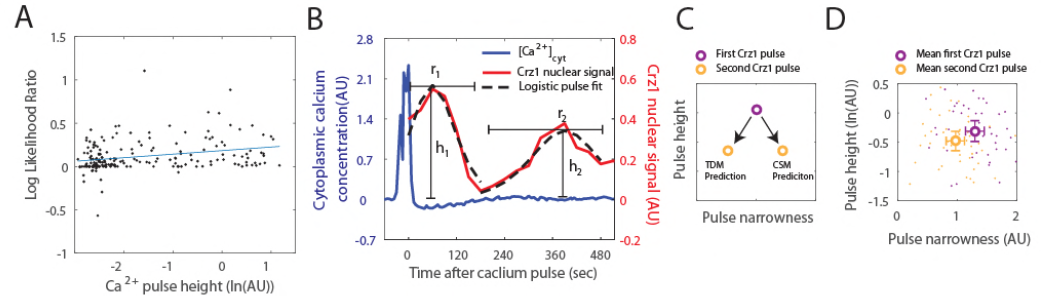
Predictions of the time delay model are confirmed by experimental data. A) Log likelihood ratio between a periodic and an aperiodic Gaussian process model shows an increasing preference towards the periodic process when calcium pulse height increases (blue line shows a linear fit, R^2^ = 0.07). Each dot represents the Crs1 trajectory following a calcium pulse. B) shows an example of logistic pulse fit (dotted trace) to experimental data (red trace) for estimating the narrowness (r_1_ and r_2_) and the heights (h_1_ and h_2_) of the first and second Crz1 pulses after a calcium pulse. C) Conformational switch model (CSM) predicts that the second Crz1 pulse is shorter and narrower than the first Crz1 pulse, while the time delay model (TDM) predicts that the second Crz1 pulse is shorter and less narrow than the first. D) Unfilled circles indicate the mean height and narrowness of Crz1 pulses. The first and second Crz1 pulses are purple and yellow, respectively. Each dot corresponds to a single Crz1 pulse identified following a calcium pulse.

Additional predictions of the time delay model are that, after a calcium pulse that is followed by at least two Crz1 pulses, the second Crz1 pulse is shorter and wider than the first Crz1 pulse because the coordinated transport of Crz1 molecules disperses across time. Although both the conformational switch model and the time delay model predict shorter second Crz1 pulses, the conformational switch model would predict a narrower second Crz1 pulse as the calcium oscillations decay (Figure 5C). To test these predictions, we identified the calcium pulses that are followed by two Crz1 pulses, and fit them to a “logistic pulse model” (Figure 5B, median R^2^ = 0.85, mean R^2^ = 0.79, See Methods) to estimate pulse height and narrowness. We found that the height of the second pulse is significantly smaller (Figure 5D, paired t-test, p < 0.005, n =42), and that the mean narrowness of the second pulses is significantly smaller than that of the first pulses (Figure 5D, paired t-test, p < 10^−8^, n = 42), supporting the time delay model to the exclusion of the conformational switch model.

Thus, the data support three additional predictions of a simple stochastic model of Crz1 nuclear import and export. Together with the explanation of the analog to digital converter, our results support the idea that coordination of Crz1 localization (and thus pulsatility) is the result of a time-delay in nuclear import and export (see Discussion).

## Discussion

Our results show that [Ca^2^+]_cyt_ is linked to Crz1 pulsatility through an analog to digital conversion: larger calcium pulses lead to more Crz1 pulses. Through this model, we can explain Crz1 pulses found after calcium pulses and provide a possible link between irregular Ca^2^+ oscillation and transcription(17). However, in our cells, Crz1 fluctuations are also found without preceding calcium pulses. These Crz1 fluctuations are aperiodic and do not show correlations between number of pulses and calcium size. We recorded movies at four different [Ca^2^+]_ext_, and found that, although the frequency of calcium pulses is correlated with external calcium concentration, the increase in calcium pulsing frequency is not comparable to the increase in Crz1 pulsing frequency, and the average size of calcium pulses does not increase significantly. Therefore, our model cannot fully explain the calcium concentration dependence of Crz1 fluctuations. However, cross-correlation analysis shows that these Crz1 dynamics have only a small correlation with [Ca^2+^]cyt dynamics (supplementary figure 1). We suggest that other intrinsic environmental fluctuations that affect Crz1 localization, such as light-(32), osmotic pressure-(33), or glucose-(34,35) induced Crz1 regulation might be involved in producing these fluctuations.

One of the interesting properties of pulsatile dynamics is that the pulses in individual cells are not synchronized, despite cells experiencing the same environmental stress(36). The analogdigital converter model explains this aspect of Crz1 pulsatility by arguing that the [Ca^2+^]cyt among individual cells at a given time point is stochastic, perhaps due to spontaneous Ca^2+^ transients(37). This model predicts that, if calcium pulses among cells could be synchronized in time, then Crz1 pulsatility should be synchronized immediately after, and that this induced synchrony would gradually decay. Consistent with this, in our nifedipine treated cells where large calcium pulses were induced in every cell immediately after calcium was added to the media, the Crz1 pulses following these synchronized calcium pulses are also synchronized, and this synchrony decays with time (Figure 3B). However, by simulating Crz1 dynamic many times with identical parameter values (Table 1), we found that our stochastic time delay model does not predict this loss of synchrony at the population level. This suggests that additional sources of cell-to-cell variability are likely missing from the stochastic time delay model.

Nevertheless, our stochastic time delay model has several advantages over a conformational switch model that assumes Crz1 nuclear localization sensitively reads out [Ca^2+^]cyt when [Ca^2+^]cyt passes through a threshold(22). Once a damped calcium oscillation is present, the conformational switch model can generate Crz1 pulses as a readout of [Ca^2^+]_cyt_ passing through a threshold (supplementary figure 4). Although we do not observe calcium oscillations passing a threshold in our movies, it is possible that our calcium sensor is not sensitive enough to distinguish these dynamics from background noise. Both our model and the conformational switch model require no negative feedback in the calmodulin/calcineurin signaling pathway. However, our model predicts the width of the second Crz1 pulse to be wider than the first, while the conformational switch model predicts the opposite: a narrower second Crz1 pulse because it is reading out a smaller fluctuation in [Ca^2^+]_cyt_. The comparison of pulse widths (Figure 5D) supports the time delay model. We also note that the stochastic model is simpler (fewer parameters needed to generate pulses and no assumption a sensitive threshold), and can directly explain the coordination of the subcellular localization of the ˜500 Crz1 molecules in the cell through time-delay in nuclear transport.

One crucial assumption in our model for the coordination among Crz1 molecules is the deactivation rate of calcineurin. Previous studies show that calcineurin has a deactivation rate *in vitro* of 0.08 fold change per minute while both calcium ions and calmodulin are presented, and has an even slower deactivation rate when either of them is not presented(38-40). The deactivation rate is slow enough to maintain the synchronous translocation of each Crz1 molecule in our model, which only requires calcineurin to return to baseline activity around 5 minutes after a calcium pulse, a length of time that has been reported *in* vitro(38-40). A conclusive test of our model would be a mutation in calcineurin that solely affects the deactivation rate, but no such mutant is available to our knowledge.

Previous work on Crz1 pulsatility suggested that Crz1 pulses are actively generated rather than passively reading out the fluctuation in [Ca^2^+]_cyt_(9). If our model is correct, then it suggests a third possibility: individual Crz1 molecules read out [Ca^2+^]cyt with a time delay. This possibility can explain the observation that higher affinity of calcineurin docking site on Crz1 leads to higher pulsing frequency(9), because higher affinity allows Crz1 to be dephosphorylated by a lower fraction of activated calcineurin and, therefore, oscillate longer after a calcium pulse. The time delay is assumed to be created by the transport between cytoplasm and nucleus, which because it requires a complicated series of steps, leads to a transport rate in the order of minutes(41). This model can be generalized to relocalization of other pulsatile transcription factors and macromolecules that have dynamics on the order of minutes, and can explain how signals that are short and fluctuating are converted into the frequency of pulses without a negative feedback loop.

## Materials and methods

### Yeast cell strain and growth conditions

BY4741 was used to construct the dual Crz1-Calcium reporter strain. Plasmids expressing GCaMP3 calcium reporter were constructed using Gibson assembly protocol(42) and gel purification. The calcium reporter gene was assembled between the promoter of ribosomal protein L39, RPL39, and the ADH1 terminator. pRPL39-GCaMP3 -tADH 1 was integrated at the *HO* locus using a selectable marker (LEU2) and confirmed by Sanger sequencing. Four replicates were performed and all showed expected GCaMP3 expression(15,16). To tag Crz1 with mCherry at the C terminus, genomic integration of pCrz1-ymCherry was done at the *CRZ1* locus using a selectable marker (URA3) and confirmed by PCR. All transformations were performed using the standard lithium acetate procedure(43).

All the time-lapse imaging experiments were started when cells were in log-phase (4 hours after being diluted from overnight liquid culture). Cells were grown in synthetic complete (SC) media lacking leucine and uracil to maintain section of markers. Carbon source was 2% glucose. For artificially increased calcium pulse experiments, 200μπι Nifedipine were added during the 4 hour inoculation.

### Spinning-disk Confocal Microscopy and image analysis

Nikon CSU-X1 was utilized for time-lapse imaging at room temperature (22° C). For GCaMP3, 488 nm laser was applied with time resolutions of 6 sec/frame, exposure time of 100 msec, and 25% laser intensity; for mCherry, 561 nm laser was applied with time resolution 30 sec/frame, exposure time of 700 msec, and 50% laser intensity. Bright field images with out-of-focus black cell edge were acquired every minute for cell segmentation and tracking.

Cells were attached to glass-bottom dishes with 0.1 mg/ml Concanavalin-A as a binding agent using a standard protocol(44,45). For each experiment, a time-lapse image series without calcium stress induction was recorded as a negative control. At the beginning of each time-lapse image series, an area of the dish that had not been exposed to laser was recorded in order to avoid blue light stress, which is known to induce Crz1 nuclear localization(32). Calcium chloride solution was added to the dish to a final concentration of 0.2M through a syringe within 20 seconds. To record the dynamics during steady state, time-lapse movies of 30 min or 1 hour were recorded after more than 1 hour of calcium stress induction for two to four time-lapse movies in each experiment. 27 replicates of time-lapse movies (18 hours in total) were recorded. Every analysis was done in both 1 hour and 30 minute time-lapse experiments.

Segmentation was automatically performed by identifying the area within cell edge through MATLAB Image Segmentation Toolbox, and cell tracking was performed by identifying 90% overlapping cell areas between two time frames. Mis-segmented and mis-stracked objects were manually removed. 23-87 cells were identified in each time-lapse movie. Single cell photobleaching correction was conducted after single cell reporter intensities were quantified (see below) using bi-exponential regression(46): for GCaMP3 intensity, correction was performed according to baseline intensity; for Crz1 expected nuclear signal, correction was performed according to Crz1 expected cytoplasmic signal. Baseline was normalized to 0 after photobleaching correction.

### Osmotic shock reduction

Crz1 localizes into the nucleus for 10 to 15 minutes after an osmotic shock (33). In the experiments where we artificially increased calcium pulses (nifedipine treatment), the effect from osmotic shock was undesirable because the calcium pulses occur immediately after addition of calcium. Change in osmotic pressure due to 0.2M calcium chloride is around 3.4 Pa, so prior to the experiment, sodium chloride solution was added (to reach 0.4M) to increase osmotic pressure to 6.1 Pa. When calcium chloride solution was added (so final concentrations of both sodium chloride and calcium chloride were 0.2M), the final osmotic pressure was now around 6.5 Pa, reducing the change in osmotic pressure before and after addition of calcium to around 0.5 Pa.

### Reporter intensity quantification

GCaMP3 intensity for each time point was estimated as average pixel intensity for all pixels in the cell.

Nuclear localization for each time point was quantified by fitting a mixture of a Gaussian distribution and a uniform distribution, and the parameters of distributions were estimated using expectation-maximization on the pixel data from each cell (see supplementary text for more details and derivation of the algorithm).

### Peak finding, pulse analyses and periodicity

Local maxima/minima were identified with Matlab function findpeaks. To smooth fluctuations shorter than 4 time points, Savitzky-Golay filtering was applied on each trajectory before defining the Crz1 pulse threshold, identifying Crz1 pulses, and quantifying calcium overshoot depth

To define the threshold for pulses, every local maximum in all cells growing in standard liquid culture (no additional calcium) was identified with a minimum distance of 60 seconds. Thresholds were then chosen to filter out most of the background noise: we chose the top 0.5% of the peak height (0.09) for calcium pulses, and the top 5% of both the peak height (0.30) and prominence (0.15) for Crz1 pulses.

For the analysis of the relationship between calcium pulse height and number of Crz1 pulses, the Crz1 pulses following a calcium pulse were counted until the next calcium pulse or the end of the time series, and the Crz1 pulses before a calcium pulse were counted until the previous calcium pulse or the beginning of the time series.

For every cell that has its largest calcium pulse after 5 minutes from the beginning or before 5 minutes from the end of the time-lapse experiments, its Crz1 trajectory was separated into pre-calcium-pulse and post-calcium-pulse trajectories. Each part was quantified if a trajectory prefers an aperiodic Gaussian process model or a periodic Gaussian process model with log likelihood ratio (LLR) using established method and MATLAB scripts(47).

### Logistic pulse fitting

A least squares method is developed to quantify Crz1 pulse height and width based on the analytic solution of logistic curve (see supplementary text for more details and derivation).

#### Acknowledgements

We thank Drs. M. Woodin, M. Cyert, M. Elowitz, N. Madras and T. Perkins for discussions. We thank Drs. M. Cyert, M. Elowitz, J. Garcia-Ojalvo, and P. Swain, as well as R. Martinez-Corral and members of the Moses lab for comments on the manuscript. This research was supported by an NSERC discovery grant to AMM and infrastructure obtained with grants from the Canada Foundation for Innovation to AMM and SP.

## Author contributions

I. S. H. performed the experiments. B. S. and S. P. provided experimental training and support. I. S. H. and A. M. developed the computational models. A. M. supervised the project. I. S. H. and A. M. wrote the paper.

